# Activation of TrkB in Parvalbumin interneurons is required for the promotion of reversal learning in spatial and fear memory by antidepressants

**DOI:** 10.1101/2022.09.07.506934

**Authors:** Elias Jetsonen, Giuliano Didio, Frederike Winkel, Maria Llach Pou, Chloe Boj, Laura Kuczynski-Noyau, Vootele Võikar, Ramon Guirado, Tomi Taira, Sari E. Lauri, Eero Castren, Juzoh Umemori

## Abstract

Critical period-like plasticity (iPlasticity) can be reinstated in the adult brain by several interventions, including drugs and optogenetic modifications. We have demonstrated that a combination of iPlasticity with optimal training improves behaviors related to neuropsychiatric disorders. In this context, the activation of TrkB, a receptor for BDNF, in Parvalbumin positive (PV^+^) interneurons has a pivotal role in cortical network changes. However, it is unknown if the activation of TrkB in PV^+^ interneurons is important for other plasticity-related behaviors, especially for learning and memory. Here, using mice with heterozygous conditional TrkB deletion in PV^+^ interneurons (PV-TrkB hCKO) in Intellicage and fear erasure paradigms, we show that chronic treatment with fluoxetine, a widely prescribed antidepressant drug that is known to promote the activation of TrkB, enhances behavioral flexibility in spatial and fear memory, largely depending on the expression of the TrkB receptor in PV^+^ interneurons. In addition, hippocampal long-term potentiation (LTP) was enhanced by chronic treatment with fluoxetine in wild-type mice, but not in PV-TrkB hCKO mice. Transcriptomic analysis of PV^+^ interneurons after fluoxetine treatment indicated intrinsic changes in synaptic formation and downregulation of enzymes involved in perineuronal net (PNN) formation. Consistently, immunohistochemistry has shown that the fluoxetine treatment alters PV expression and reduces PNN_s_ in PV^+^ interneurons, and here we show that TrkB expression in PV^+^ interneurons is required for these effects. Together, our results provide molecular and network mechanisms for the induction of critical period-like plasticity in adulthood.

## Introduction

Learning and memory dysfunction is a common neuropsychological symptom of neuropsychiatric and neurological diseases. It has been proposed that chronic treatment with antidepressants (ADs) improves impaired learning and memory in animal models [1–3] via increased neuronal plasticity, by promoting neurogenesis [4,5], and long-term potentiation (LTP) in the hippocampus [6–8]. However, it is still not clear how AD treatments improve the dysfunction.

The activation of the brain derived neurotrophic factor (BDNF) and its receptor TrkB is a key factor in neuronal plasticity. The binding of BDNF to TrkB causes the autophosphorylation of TrkB and leads to the activation of intracellular signalling pathways involved in neuronal differentiation, survival, and growth, as well as synaptic plasticity in neurons [9,10]. This pathway also regulates gene transcription and long-term potentiation (LTP) [9]. Previous studies demonstrated that chronic treatment with AD, such as the selective serotonin reuptake inhibitor (SSRI) fluoxetine, increases the plastic state of Parvalbumin-positive (PV^+^) fast-spiking interneurons primarily targeting the perisomatic area of pyramidal neurons [11–13]. Donato et al. showed that PV^+^ fast-spiking basket cells exhibit plasticity by dynamically changing their states in response to recent experience: a state characterized by low PV expression in the PV^+^ interneurons is involved in plastic networks while a state with high PV-expression in PV cells promotes memory consolidation in the hippocampal CA3 region. This leads to a lower and a higher number of excitatory synaptic inputs onto PV interneurons, respectively, regulating experience-dependent network plasticity [14]. Furthermore, perineuronal nets (PNN) [15], an extracellular matrix surrounding PV interneurons is known to be a plastic structure regulated by iPlasticity in the amygdala, hippocampus and visual cortex [13,16,17].

We have demonstrated that ADs induce a critical period-like plasticity in the adult brain (iPlasticity), which allows brain networks to better adapt to environmental stimuli, such as training or rehabilitation, and consequently ameliorate neuropsychiatric symptoms [7,10,18]. iPlasticity occurs in a variety of brain areas and can be induced by different interventions to modulate behaviors when combined with appropriate trainings. We have proposed the “network hypothesis” of neuropsychiatric diseases, according to which neuropsychiatric diseases reflect malfunctioning information processing within particular neural networks, and interventions, including ADs, act by providing an opportunity for neuronal activity to improve this processing [19]. Our laboratory recently demonstrated that ADs directly bind to TrkB through a lipid binding motif and activate TrkB to promote neural plasticity [20]. We also recently showed that TrkB activation in PV^+^ interneurons is necessary and sufficient for iPlasticity in the visual cortex [16]. Therefore, the treatment with ADs is a good tool to directly activate TrkB to study the mechanisms of iPlasticity. However, it is still unknown whether elevated plasticity by ADs in PV^+^ interneurons combined with learning processes can improve learning and memory more generally.

In order to assess the effects of TrkB in PV^+^ interneurons on reversal learning, we treated PV^+^ interneuron-specific heterozygous TrkB knockout (PV-TrkB hCKO) mice with fluoxetine in the fear extinction paradigm and in IntelliCage apparatus. We also examined the dependency of LTP on the expression of TrkB in PV^+^ interneurons by studying local field potential activity in the hippocampal CA1 of PV-TrkB hCKO mice. We then performed a transcriptomic analysis specifically for PV^+^ interneurons using translating ribosome affinity purification (TRAP) after a chronic treatment with fluoxetine, and found intrinsic changes in PV^+^ interneurons, especially in genes related to the formation of PNN_s_. Finally, we immunohistologically confirmed the plastic changes in PV^+^ interneurons after chronic treatment with fluoxetine.

## Material and Method

Details of Material and Methods are in the Supplementary Materials and Methods

### Animals and experimental design

Heterozygous mice with reduced expression of TrkB specifically in PV^+^ interneurons (PV-TrkB hCKO; PV^pvr/wt^, TrkB^flx/wt^) were produced by mating females from an heterozygous PV specific Cre line [21] (PV^pvr/wt^; Pvalb-IRES-Cre, JAX: 008069, Jackson laboratory) with males from an homozygous floxed TrkB mouse line (TrkB^flx/flx^) [22] (Fig. 1a). Due to frequent fights among males, only females (5 months old) were used for IntelliCage and the males (2 months old) were used for the fear extinction paradigm. Transgenic mice harboring FLEX-L4 conjugating GFP [23] were crossed with homozygous PV specific Cre mice (PV^pvr/wt^) to obtain the mice (2 months old) expressing GFP-L4 specifically in PV interneurons. The room temperature was kept at 23±2°C, and all mice were kept in a room with a 12-hr light/dark cycle (lights on at 6:00 a.m.) with access to food and water *ad libitum*. All experiments were carried out in accordance with the European Communities Council Directive 86/6609/EEC and the guidelines of the Society for Neuroscience and were approved by the County Administrative Board of Southern Finland (License number: ESAVI/38503/2019).

**Fig. 1.**
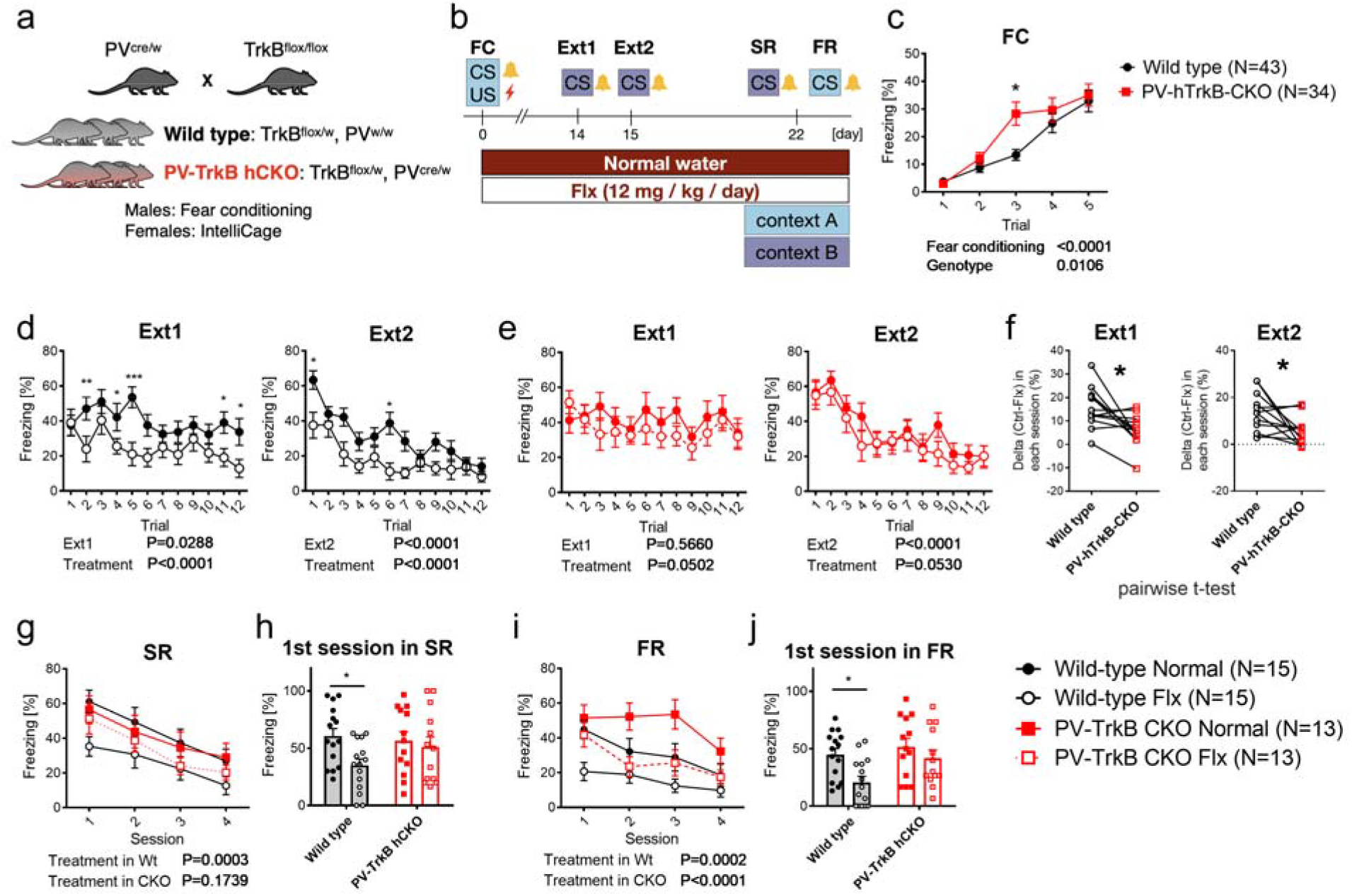
Fluoxetine treatment promotes contextual and cued fear erasure and depends on TrkB expression in PV interneurons. (a) Mating strategy to obtain wild type and PV specific heterozygous TrkB knockout mice PV-TrkB hCKO). (b). Scheme of the fear conditioning paradigm. Mice were conditioned by pairing a tone and an electric shock in context A (c), and one group was treated with fluoxetine (24mg/kg/day). After two weeks, mice were subjected to two days of fear extinction training: day1 (Ext1), day 2 (Ext2) *wt* (d), PV-TrkB hCKO (e). After one week, mice were tested for spontaneous recovery (SR) in context B (g, h) and fear renewal (FR) in context A (i, j). (c) Both wild-type and PV-TrkB hCKO mice increased freezing during the conditioning/acquisition phase and exhibited the same levels of fear acquisition (two-way ANOVA, Conditioning, F (4, 375) = 33.81, P < 0.0001). However, PV-TrkB hCKO mice showed significantly higher freezing compared to wild-type mice (Genotype, F (1, 375) = 6.601, P < 0.0106; post-hoc, wild-type vs PV-TrkB hCKO in Trial 3, p=0.0186). (d) In wild-type mice, significant effects of sessions were detected in Ext1 and Ext2 (Extinction, Ext1, F (11, 348) = 1.869, P = 0.0423; Ext2, F (11, 348) = 9.655, P < 0.0001) and an effect of fluoxetine treatment on both days (Treatment, Ext1, F (1, 348) = 36.74, P < 0.0001; Ext2, F (1, 348) = 44.24, P < 0.0001). (e) PV-TrkB hCKO mice tended to have an effect of fluoxetine treatment (2-way ANOVA, treatment, Ext1, F (1, 288) = 3.866, P = 0.052; Ext2, F (1, 288) = 3.766, P = 0.053), and the extinction training significantly reduced the freezing in all PV-TrkB-hCKO mice only on day 2 (Ext2, session, F (11, 288) = 9.655, P < 0.0001). (f) Pairwise t-test of fluoxetine effect (delta: freezing in control (%) – fluoxetine (%)) between wild-type and PV-TrkB hCKO mice on each session in two days. Wild-type mice had a significantly higher difference compared to PV-TrkB hCKO mice on sessions in Ext1 (paired t-test, p = 0.0101) and Ext 2 (paired t-test, p = 0.0228). (g) In SR, fluoxetine treatment significantly decreased freezing throughout sessions in wild-type mice (2-way ANOVA, Treatment, F (3, 116) = 15.15, p = 0.0002) but not in PV-TrkB hCKO mice (2-way ANOVA, treatment, F (1, 96) = 1.876, p = 0.1739) (h). In the 1^st^ session of SR, fluoxetine treatment decreased freezing in the wild type mice (Holm-Sidak’s multiple comparisons test, p = 0.0229) but not in PV-TrkB hCKO mice (p = 0.6282). (i) In FR, fluoxetine treatment induced a decrease of freezing in both wild-type (2-way ANOVA, treatment, F (1, 116) = 10.14, P = 0.0020) and PV-TrkB hCKO mice (treatment, F (1, 96) = 16.61, p < 0.0001). (j). In the 1^st^ session of FR, fluoxetine treatment decreased freezing in wild type (Holm-Sidak’s multiple comparisons test, p = 0.0122) but not PV-TrkB hCKO mice (p = 0.2892). Error bars designate SEM. *p < 0.05; **p < 0.01.

### Fear extinction paradigm

The fear conditioning paradigm was conducted following a protocol described previously [13].

### IntelliCage with chronic fluoxetine treatment

Intellicage (NewBehavior AG, Zurich, Switzerland) is an automated device that allows housing, performance and measurements of specific tasks in a fully automated manner, removing the need for a human operator and operator-derived bias [24,25]. Mice were divided into two groups: the “control” treated with 0.1% (w/v) saccharine in the drinking water and the “experimental group” with 0.1% (w/v) saccharin supplemented with 0.08% (w/v) fluoxetine in the drinking water. We performed patrolling tasks, where the water bottles were made accessible (doors would open for 4s) only if the mouse nose-pocked the “active” door area which, once discovered and used by the mouse, would switch to the one immediately next to it, in a clockwise direction. During the Reversal phase the direction was switched to counter-clockwise.

### Electrophysiology in acute slices

Field excitatory postsynaptic currents (fEPSPs) were recorded in an interface chamber using ACSF-filled electrodes (2-4 MΩ) positioned within the CA1 stratum radiatum (Supplemental note). Notably, LTP was induced through tetanic stimulation (200ms pulse interval; 100 pulses; 0.1ms pulse duration) and recorded for 45min.

### Immunohistochemistry

Animals treated with control or fluoxetine were perfused transcardially with PBS followed by 4% PFA in PBS and the brains were isolated. The brains were post-fixed overnight and stored in PBS with 0.02% NaN_3_ until cutting on a vibratome (VT 1000E, Leica). Free-floating sections (40 μm) were processed for fluorescence immunohistochemistry following a protocol described previously [26] using antibodies listed in supplemental table 1.

### Confocal imaging and imaging analysis on PV and PNN intensity

Immunohistologically stained sections were imaged with a confocal microscope (Zeiss LSM 700). PV/PNN was imaged in different sections containing CA3b region in the dorsal hippocampus (between −1.94 and −2.18 mm in the Anterior-Posterior axis relative to Bregma). The multilayer confocal images were stacked (z-stack, maximum intensity), and the fluorescence intensity of PV, PNN and TdTomato were analyzed with Fiji software (National Institute of Health, US) (https://fiji.sc/) [27]

### TRAP sample preparation and sequencing

TRAP-analysis was performed according to a previously published protocol [28]

### Experimental design and statistical analysis

Biochemical and behavioral data were analyzed by two-way ANOVA, taking sessions, genotype, and treatment with fluoxetine, followed by Fisher’s LSD test. All results of two-way ANOVA are shown in the supplemental table 3. All statistical analyses were performed using Prism 6 or 8 (GraphPad Software). A *p*-value <0.05 was considered statistically significant.

## Results

### Expression of TrkB in PV+ interneurons is important for fear erasure induced by fluoxetine treatment

We have previously demonstrated that a chronic treatment with fluoxetine combined with fear extinction training promotes the erasure of previously acquired fear memory and alteres the configuration of PV^+^ interneurons [13]. We first tested whether this promoted fear erasure might depend on TrkB expressed in PV^+^ interneurons and would therefore be blunted in PV-TrkB hCKO mice (Fig. 1a). In a fear conditioning paradigm (Fig. 1b), all mice were conditioned with a shock paired with a sound cue in context A during the fear-conditioning/acquisition phase, resulting in an increased freezing that was comparable in duration across all groups, although PV-TrkB hCKO mice conditioned faster than wild-type mice (Fig. 1c). The control and PV-TrkB hCKO mice were then assigned equally and randomly into groups receiving either water or water supplied with 0.08% (w/v) of fluoxetine, both enriched with 0.1 % (w/v) saccharin. Two weeks later, the mice were exposed to the conditioned stimulus (CS; “beep” sound) in context B during 2 days of extinction training. In the wild-type group, both control and fluoxetine treated mice showed a decreased freezing, but the effect was significantly more pronounced in the fluoxetine treated group (Fig. 1d). Both control and fluoxetine treated PV-TrkB hCKO mice showed decreased freezing on the 2^nd^ day of extinction training, but the response to fluoxetine was significantly less pronounced than in the wild-type mice (Fig. 1e). In fact, when comparing the difference in freezing (delta) between water-treated and fluoxetine-treated mice in each session, the wild-type mice showed a significantly higher difference compared to PV-TrkB hCKO mice by pairwise t-test (Fig. 1f), suggesting that in the absence of TrkB in PV neurons, the effects of fluoxetine are significantly reduced. One week later, the fluoxetine-treated wild-type mice showed decreased freezing throughout the whole session in context B (spontaneous recovery, SR) (Fig. 1g), as well as in the 1^st^ session of this test (Fig. 1h). However, the PV-TrkB hCKO mice failed to show similar effects of the fluoxetine treatment throughout the sessions of the test (Fig. 1g, h). In addition, the treatment significantly reduced the freezing in the fear renewal test (FR) in context A in wild-type mice, especially in the first session of (Fig. 1i, j). Interestingly, the treatment with fluoxetine decreased the overall freezing of PV-TrkB hCKO mice in the fear renewal test (Fig. 1i), but there was no difference in the first session (Fig. 1j). These results suggest a role of TrkB expression in PV neurons in the extinction-enhancing effects of fluoxetine in cued fear conditioning, but a less pronounced role in the contextual component of the paradigm (FR).

### Expression of TrkB in PV+ interneurons is important for the improvement of reversal spatial learning induced by fluoxetine treatment

The IntelliCage experiments were conducted to test the effect of chronic fluoxetine treatment on spatial learning as depicted in Fig. 2a. Mice were implanted with transponders and were treated with fluoxetine-containing water for two weeks before the experiments. During the adaptation to freely accessible water bottles in the corners (FA), nose pokes (NPA), and drinking sessions (DSA), six mice were excluded because they could not learn the adaptation tasks [Control group (wild-type, 1; PV-TrkB hCKO, 2), fluoxetine-treated group (wild-type, 1; PV-TrkB hCKO, 2)]. In the acquisition phase of the patrolling task, the location of the open corner changed after each visit, and the water-deprived mice had to patrol the corners in a “clockwise” order to receive a water reward (Fig. 2a, left panel). The percentages of error ratios were calculated as the number of visits in the incorrect corner divided by the number of total visits. The wild-type mice decreased the error ratio during sessions, and there was no effect of fluoxetine treatment in the acquisition phase (Fig. 2c-e). The PV TrkB hCKO mice also decreased the error ratio during sessions (Fig. 2f-h), but interestingly the fluoxetine treatment decreased the error ratio faster than in water-treated mice (Fig. 2f). The PV TrkB hCKO mice had significantly higher error ratios compared to wild-type mice treated with control water (Supplemental fig 1a). These results indicate that PV TrkB hCKO mice have lower spatial learning skills in acquisition compared to wild-type mice, but the fluoxetine treatment recovers them to a level comparable to wild-type mice.

**Figure 2.**
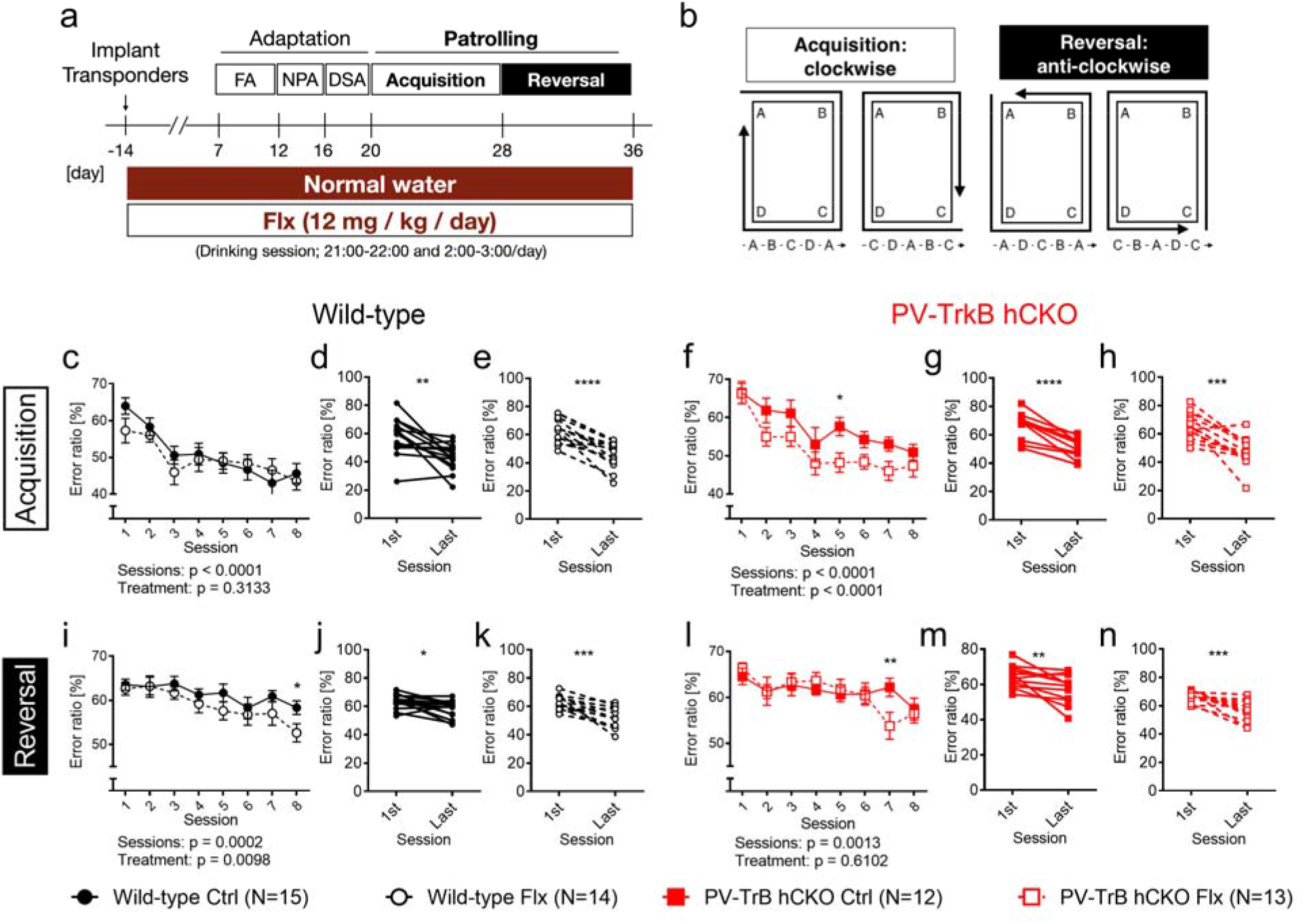
Chronic treatment with fluoxetine promotes spatial learning and depends on TrkB expression in PV interneurons. (a) Scheme of the Intellicage system during the chronic treatment with fluoxetine. All mice were implanted with transponders, and were treated with fluoxetine in water. Mice adapted gradually to the tasks in the IntelliCage (Free adaptation, FA; Nose poke adaptation, NPA; Drinking session adaptation, DSA), followed by the actual leaning tasks (Patrolling). (b) Scheme of the patrolling task. Error ratio in acquisition (c-h) and reversal phase (i-n) in wild-type (c-e, i-k) and PV-TrkB-hCKO mice (f-h, l-n) (n = 12– 15 per group). (c) Wild-type mice decreased the error ratio during the sessions (sessions, F (7,128) = 4.905, p < 0.0001), but there was no difference caused by the treatment (two-way ANOVA, treatment, F (1, 128) = 2.077, p = 0.3133). Significant differences were found in pair-wise comparisons between the 1^st^ and last sessions in control mice (pairwise t-test, p = 0.0031) (d) and fluoxetine water-treated mice (p < 0.0001). (e). Fluoxetine treatment reduced the error ratio during sessions in PV-TrkB-hCKO mice (treatment, F (1, 183) = 16.37, p < 0.0001; sessions, F (7, 183) = 9.462, p < 0.0001). There was a significant difference in the error ratio between the initial and the last sessions in control (pairwise t-test, p < 0.0001) (g) and fluoxetine-(p = 0.0001) (h) treated mice. (i) In the reversal phase, wild-type mice showed a significant effect in both sessions (F (7, 208) = 4.212, p < 0.0002) and treatment (F (1, 128) = 2.077, P = 0.0098). There were significant differences between the error ratio in the initial session and the last one in both control (pairwise t-test, p=0.0097) (h) and fluoxetine treated mice (p=0.0006) (i). (l) In PV hTrkB cKO mice, fluoxetine treatment did not have an effect during sessions (F (1, 184) = 0.2608, p = 0.6102). There was a significant difference in error ratios between 1^st^ and last session in control (pairwise t-test, p = 0.0097) (m) and fluoxetine treatment (p = 0.0006) (n). Error bars designate SEM. *p < 0.05; **p < 0.01.

In the reversal phase, wild-type mice significantly reduced the error ratio during sessions in both control and fluoxetine-treated groups (Fig. 2i-k), but the treatment with fluoxetine facilitated the decrease of the error ratio during sessions, especially in the final session (Fig. 2i). These results indicate that the fluoxetine treatment improves the reversal learning in wild-type mice. The PV hTrkB CKO mice also improved their performance during sessions (Fig. 3l-n). Notably however, there was no effect of the fluoxetine treatment (Fig. 3l). These results strongly suggest that TrkB expression in PV interneurons is important for the effect of fluoxetine on the reversal learning in spatial tasks.

**Fig. 3.**
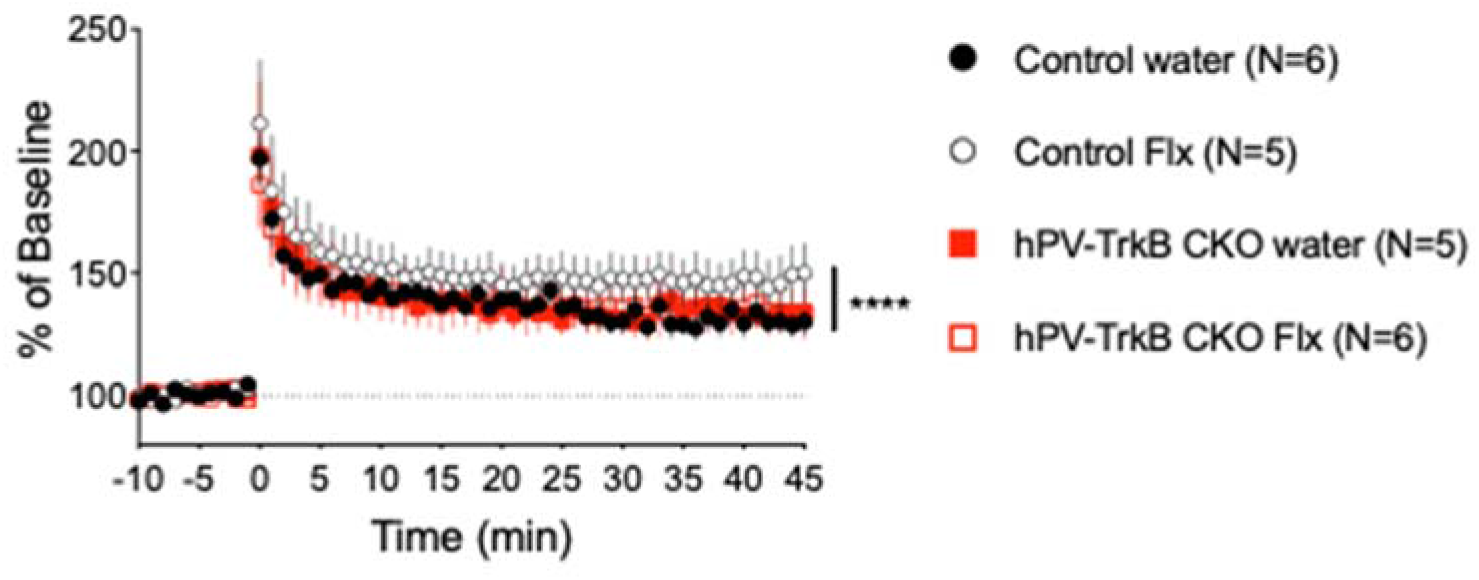
Chronic treatment with fluoxetine enhances synaptic plasticity in the hippocampus. LTP induction after chronic treatment with fluoxetine. LTP was significantly enhanced 45 min after tetanic stimulation in wild-type mice treated with fluoxetine compared to control (Two-way ANOVA, treatment, wild-type, F (1, 414) = 50.20, P < 0.0001) but not in PV-TrkB hCKO mice (treatment, F (1, 491) = 0.2324, p = 0.6300). Bars indicate mean ± SEM.

### Fluoxetine treatment potentiates hippocampal LTP through expression of TrkB in PV interneurons

In order to understand whether the improved behavioral flexibility after fluoxetine treatment reflects enhanced neural plasticity in the hippocampus, the main region involved in contextual fear and spatial memory [29], we recorded fEPSPs in acute hippocampal slices of wild-type and PV TrkB hCKO mice after chronic fluoxetine treatment (Fig. 3). As previously reported [8,30] we observed a significant enhancement of long-term potentiation (LTP) at 45 min after tetanic stimulation in wild-type mice treated with fluoxetine compared to mice treated with water. There was, however, no effect of fluoxetine treatment on LTP in hPV-TrkB CKO mice (Fig. 3). These results indicate that the chronic treatment with fluoxetine enhances LTP induction and expression, which seems to require TrkB expression in PV cells.

### PV-specific transcriptomic analysis through TRAP

In order to investigate gene expression in PV interneurons after chronic treatment with fluoxetine, we conducted a TRAP analysis to investigate ongoing protein translation specifically in PV^+^ interneurons (Fig 4a). After chronic treatment with fluoxetine, the whole hippocampus of mice expressing EGFP-tagged L10a ribosomal subunits specifically in PV interneurons (Fig. 4b) was used for TRAP followed by next generation sequencing (NGS). We found 879 genes that were differentially expressed after chronic treatment with fluoxetine (p < 0.05) and these were further studied by a pathway analysis. The chronic fluoxetine treatment significantly affected several of these pathways in the hippocampus (p < 0.1) (Supplemental table 2), and representative pathways are shown in Fig. 4c. Particularly, genes in glycosaminoglycan chondroitin sulphate and heparan biosynthesis pathways were significantly down regulated. These are associated with chondroitin sulphate proteoglycans, which are an integral part of PNN_s_ [15]. Also, genes related to glycerolipid and glycerophospholipid metabolism were down regulated. These pathways are involved in the regulation of lipid composition of the cellular membrane, which is highly related to antidepressant effects [20]. Furthermore, the fluoxetine treatment significantly changed the expression of genes in the GABAergic synapse pathway, including G Protein Alpha Inhibiting Activity Polypeptide 3 (Gnai3), G protein subunit gamma 4, 8, and 13 (Gng4, Gng8, and Gng13), which are coupled with GABA type B receptor, and mediate slow and prolonged inhibitory action [31]. Huntingtin-associated protein 1 (Hap1) directly interacts with GABA type A (GABA_A_) receptors and influences the recycling of the receptor by inhibiting its degradation [32]. Such modulation of the expression and localization of GABA_A_ receptors are thought to be a plastic event resulting in maintenance of the excitatory/inhibitory balance [33].

**Fig. 4.**
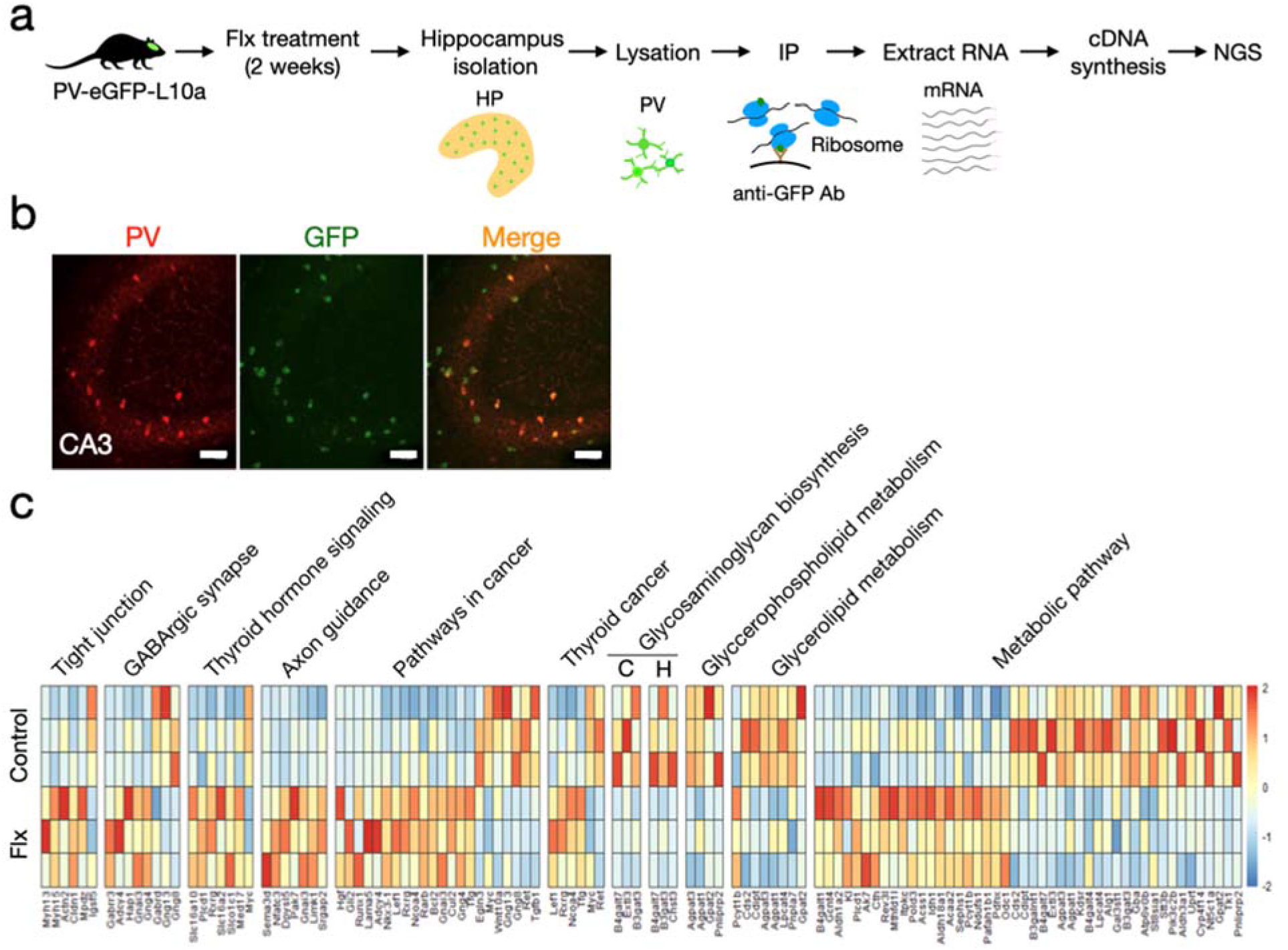
TRAP analysis of PV interneuron after chronic treatment with fluoxetine. (a) Ribosome-tagged transgenic mice were treated with fluoxetine or control water for two weeks, and their hippocampi were isolated and lysated. Ribosomes bound to mRNA were immunoprecipitated with beads coated with GFP-antibody and the mRNA was purified for cDNA synthesis followed by next generation sequencing (NGS). (b) Immunohistochemistry analysis with anti-PV antibody. Parvalbumin is co-localized with GFP indicating that the cells expressing GFP-tag in ribosomes are PV-cells. Scale bars, 50 μm. (c) Heatmap of significant genes and pathways detected by GO analysis. C, chondroitin sulfate; H, heparan sulfate.

Overall, our TRAP analysis revealed a novel mechanistic insight into the observed phenomena of increased neural plasticity in PV^+^ interneurons, such as synaptic formation and turnover of PNN_s_ through the regulation of gene expression after fluoxetine treatment.

### Decreased intensity of PV and PNN after fluoxetine treatment depending on TrkB expression in PV^+^ interneurons

TRAP analysis showed decreased expression of genes related to the formation of PNN. In addition, it has been reported that PV configurations in the CA3 region of the hippocampus are dynamically regulated by experiences, such as environmental enrichment and fear conditioning [34]. We used immunohistochemistry to analyze the intensities of PV, and PNN_s_ surrounding PV interneurons as a measure of their expression levels in the hippocampal CA3 region after chronic fluoxetine treatment (Fig. 5a). After fluoxetine treatment the proportion of low-intensity PV cells increased, and the high-intensity PV cells were reduced in wild-type mice, while there was no obvious difference in the proportions of PV intensity in PV-TrkB hCKO mice (Fig. 5b). In addition, the proportion of PV-positive cells among cells positive for PNN was significantly reduced after fluoxetine treatment in wild-type mice as shown previously [13], but not in PV TrkB hCKO mice (Fig. 5c). Interestingly, when the intensity of PNN_s_ were separately measured in lower- and higher- PV expressing PV interneurons, the fluoxetine treatment significantly reduced the intensity of PNN only in high- but not in low- PV-expressing cells in wild-type mice (Fig. 5d). However, the treatments showed no effect on the PNN intensity between in either low- or high- PV expressing cells in PV-TrkB hCKO mice (Fig. 5e). These results strongly suggest that chronic fluoxetine treatment shifts the configuration of PV interneurons towards lower PV and PNN expressing cell state through TrkB signaling. Taken together with the TRAP analysis, the decreased gene expressions of the extracellular matrix might be involved in the reduced PNN formation after chronic treatment with fluoxetine.

**Fig. 5.**
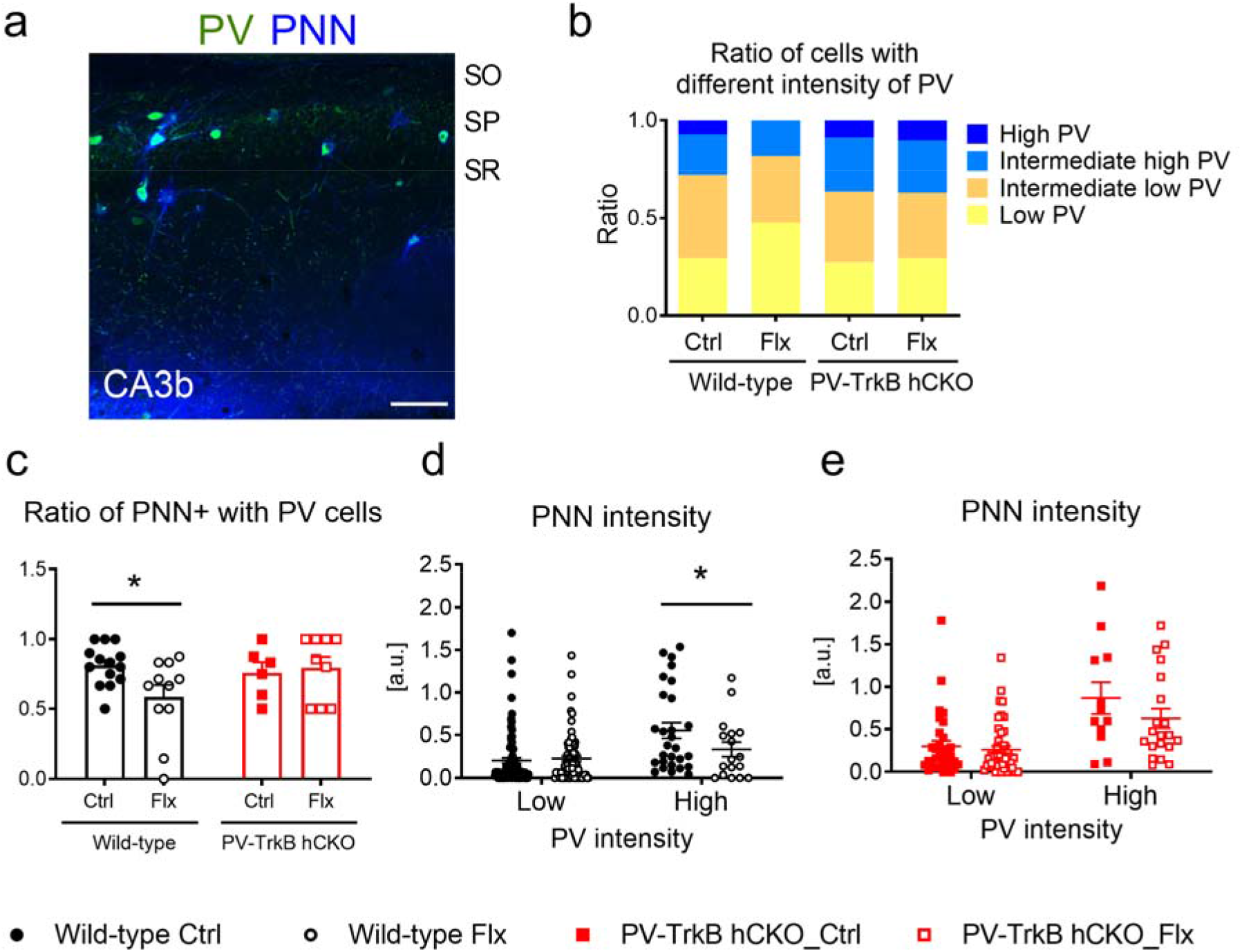
Chronic treatment with fluoxetine enhances PV plasticity in the hippocampus. (a-e) Image analysis on PV and PNN expression in the dorsal hippocampus of control and fluoxetine-treated wild type and PV-TrkB hCKO mice. (a) Representative image of PV and PNN staining. Immunostaining with PV and PNN followed by intensity analysis on PV and PNN. SP, stratum pyramidale; SO, stratum oriens; SR, stratum radiatum. Scale bar, 200 μm. (b) Intensity analysis of PV expression in PV interneurons. The ratio of high and intermediate-high PV was lower after fluoxetine treatment in wild-type mice, but this difference was not observed in PV-TrkB hCKO mice. (c) Fluoxetine-treated wild-type mice have significantly lower percentages of PV interneurons also expressing PNN_s_, but this effect is blunted in PV-TrkB hCKO mice (Fisher’s LSD post-hoc test, control vs Flx: wild-type, p = 0.0138; CKO, p = 0.7571). (d, e) PNN intensity analysis in cells separated by PV-intensity. Fluoxetine treatment reduces PNN intensities in high-(intermediate-high and high) PV expressing cells only in WT mice (two-way ANOVA, interaction between, F (1, 208) = 4.785, P= 0.0298, Fisher’s LSD post hoc test, control vs Flx: Low, p = 0.6365; High, p = 0.0281), but not in PV-TrkB hCKO mice (two-way ANOVA, interaction, F (1, 108) = 1.326, P=0.2520, Fisher’s LSD post hoc test, control vs Flx: Low, p = 0.6722; High, p = 0.1075).WT, control, low, n = 88; WT, Flx, low, n = 79; WT, control, high, n = 28; WT, co, high,17; CKO, control, low, n = 32; CKO, Flx, low, n = 48; CKO, control, high, n = 12; CKO, control, high,20. Bars indicate mean + SEM. *p < 0.05

## Discussion

Here, we demonstrate that iPlasticity induced by pharmacological activation of TrkB in PV^+^ interneurons promotes reversal learning in fear and spatial memory. We also showed that both the potentiation of LTP in Schaffer collaterals to CA1 synapses, and the shift in the configuration of the PV and PNN network to a more plastic state require TrkB expression in hippocampal PV^+^ interneurons. These alterations involve changes in the gene expression patterns related to GABAergic synapses and PNN formation.

### Impaired fear extinction consolidation of heterozygous PV-TrkB CKO mice

It has previously been shown that heterozygous PV-hTrkB CKO male mice exhibit slightly impaired extinction consolidation but not fear acquisition/conditioning in auditory fear paradigm [35]. In this study, PV-hTrkB CKO mice showed normal or even faster fear acquisition, but control heterozygous PV-TrkB CKO mice kept a higher level of freezing in fear renewal, compared to that of wild-type mice. Altogether, heterozygous PV-TrkB CKO mice seem to a have deficit in the consolidation of fear extinction.

### Roles of TrkB in PV^+^ interneurons for contextual fear erasure

We show here that fluoxetine treatment decreases cued and contextual fear responses in wild-type mice as previously shown [13], but not in heterozygous PV-TrkB CKO mice. However, fluoxetine treated heterozygous PV-TrkB CKO mice showed a blunted effect only in the initial phase, but then reduced their fear responses in later phases during fear renewal. Interestingly, optical activation of TrkB in the pyramidal neurons of the ventral hippocampus showed a similar pattern: decreased contextual fear response except for the initial phase [26]. These observations raise the possibility that the expression of TrkB in PV interneurons is important, but that TrkB expression in pyramidal neurons is also involved in the formation of a new inhibitory memory to overwrite a conditioned memory or to erase a fear memory.

### PV interneurons are involved in the reversal learning phase in spatial learning

To test whether fluoxetine treatment can improve behavioral flexibility or spatial reversal learning through TrkB receptors in PV interneurons, we used the IntelliCage system. IntelliCage is an automated setup that allows behavioral experiments without direct handling of mice except for when changing cages and water bottles, which results in higher reproducibility and reliability [36]. Previous studies have shown that CKIIa specific TrkB heterozygous CKO mice had normal spatial memory in the classical Morris water maze test, but had a lower behavioral flexibility in a naturalistic settings [22,37]. Chronic treatment with fluoxetine did not affect acquisition but it enhanced reversal spatial memory in wild-type mice. In contrast, PV-TrkB hCKO mice showed improved spatial memory in acquisition but not in reversal phase after fluoxetine treatment. In addition, PV-TrkB CKO mice showed a higher error ratio in acquisition but not in reversal phase, when compared to the wild-type mice. These observations, suggest that TrkB in PV^+^ interneurons play a central role in both the basal level of spatial learning as well as in newly formed memory, especially reversal learning promoted by fluoxetine.

### PV and PNN as makers of plasticity in PV interneurons

We found that fluoxetine treatment mainly affects the high-PV expressing PV interneurons, which have been demonstrated to be born earlier during embryonic development and to be responsible for memory formation of recent experiences [14,34]. For instance, fear conditioning increases the number of high PV-expressing cells, while environmental enrichment increases the fraction of PV basket cells with low levels of PV [34]. These observations suggest that chronic fluoxetine treatment and environmental enrichment show similar effects, promoting the more plastic state of PV configuration. In addition, we demonstrate that the intensity of PNN was reduced after fluoxetine treatment only in high PV-expressing but not in low- PV-expressing interneurons. Lower expression of PNN is known to represent a plastic state of PV interneurons [38], and it is interesting that chronic fluoxetine treatment regulates both PV and PNN_s_ via TrkB expression. The TRAP analysis also points towards the regulation of PNN_s_ as the target of fluoxetine action. We have previously shown that plasticity promoted by the reduction of PNN_s_ by chondroitinase treatment is also dependent on the expression of TrkB in PV interneurons and that this effect is mediated by the inhibition of the PNN receptor, receptor-type tyrosine-protein phosphatase Sigma (PTPRS) [39]. In addition, fluoxetine was shown to disrupt the interaction between TrkB and PTPRS, functionally mimicking the effects of PNN disruption [39]. Furthermore, it appears that fluoxetine specifically targets PV^+^ interneurons which was also observed in visual cortex plasticity [16]. These effects of fluoxetine treatment on the TrkB receptors expressed in PV interneurons might be involved in behavioral flexibility, which would promote the exchange or renewal of consolidated memories.

### LTP is increased through TrkB activation in PV interneurons

Hippocampal LTP is widely regarded as the cellular substrate underlying learning and memory, enabling plasticity processes to take place [40]. Previous research has shown that chronic fluoxetine treatment increases LTP in the hippocampus, amygdala and visual cortex [8,13,16,41]. The TrkB receptor and particularly its signaling through phospholipase Cγ (PLCγ) has emerged as a potent regulator of LTP [9]. We now show that the fluoxetine-mediated increase in hippocampal LTP is prevented when TrkB expression is reduced in PV interneurons.

### Gene regulation in fluoxetine-induced plasticity in PV^+^ interneurons

In addition to the genes related to the formation of PNN_s_, composition of cellular membrane, and regulatory proteins of GABArgic receptors, all significant DE genes related to axon guidance were up-regulated. For instance, Sem3D is a receptor of Sema3A, which is known to be localized in PNN_s_ [42], and Srgap2 is localized in synapses and regulates synaptic densities through Rac1-GAP activity[43,44].

Since the Rac1 signaling regulates the density of inhibitory synapses within dendrites and their subcellular distribution, Srgap2 is considered to coordinate excitatory/inhibitory balance [45]. These results suggest that axon regeneration and sprouting actively occurred and can potentially rewire neuronal networks involving PV^+^ interneurons responding to environmental stimuli.

### Mechanisms of fluoxetine-induced plasticity

We have recently shown that fluoxetine directly binds to the transmembrane domain of TrkB dimers and increases TrkB retention in the plasma membrane, thereby allosterically promoting BDNF signaling [20]. Furthermore, antidepressants disrupt the interaction between TrkB and the AP2 complex involved in endocytosis, promoting TrkB localization in plasma membrane [46]. Consistently with the present findings, we have observed that the activation of TrkB specifically in the PV interneurons is necessary and sufficient for iPlasticity in the visual cortex [16]. We found here that TrkB activation by fluoxetine regulates PNN_s_ encasing PV neurons and our previous findings suggest that reduction in PNN_s_ further promotes TrkB activity within PV^+^ interneurons [17]. Taken together, a positive feedback loop between reduction of PNN and TrkB activation may explain the critical role of TrkB in the PV^+^ neurons in iPlasticity. Importantly, while the activation of TrkB in pyramidal neurons promotes their excitability, in PV^+^ neurons TrkB activation reduces excitability through downregulation of Kv3-family potassium channels [16]. Therefore, the activation of TrkB in PV^+^ interneurons does not counteract the concomitant TrkB activation in pyramidal neurons but synergizes with it by disinhibiting pyramidal neurons, thereby orchestrating an enhanced state of cortical plasticity that underlies iPlasticity. Our present data suggest that a similar kind of state of enhanced plasticity, involving reformation of GABAergic signaling and reduction in PNN_s_, is underlying the effects of fluoxetine on behavioral flexibility and reversal learning.

## Acknowledgement

We thank Sulo Kolehmainen and Outi Nikkila for assistance in all experiments. We also thank the caretakers in the F-building in UH for assistance with animal care, and Sequencing unit of Institute for Molecular Medicine Finland FIMM Technology Centre, University of Helsinki supported by Biocenter Finland. Behavioural testing was carried out at the Mouse Behavioural Phenotyping Facility (MBPF) supported by Biocenter Finland and HiLIFE. We also thank Anu Suoranta,and Pirkko Mattila in FIMM Transcriptomics services supported by UH and Biocenter Finland.

## Author Contributions

JU and EC conceived of and designed the project. CB, GD, and JU performed behavioral experiments. ML, LK, and JU conducted immunohistochemistry and imaging analysis under the supervision of RG. F.W. performed electrophysiological experiments under the supervision of TT and S.L. EJ performed TRAP analysis under the supervision of JU. All authors were involved in writing the manuscript.

