## Supplemental material for "Activation of TrkB in Parvalbumin interneurons is required for the promotion of reversal learning in spatial and fear memory by antidepressants"

### Supplementary Materials and Methods

#### Animals and experimental design

Heterozygous mice with reduced expression of TrkB specifically in $PV^+$ interneurons (PV-TrkB hCKO; $PV^{pvr/wt}$, $TrkB^{flx/wt}$) were produced by mating females from an heterozygous PV specific Cre line [1] ($PV^{pvr/wt}$; Pvalb-IRES-Cre, JAX: 008069, Jackson laboratory, backcrossed with C57BL/6J for more than 10 generations) with males from an homozygous floxed TrkB mouse line ($TrkB^{flx/flx}$) [2] (Fig. 1a). Due to frequent fights among males, only females were used for the IntelliCage and the males were used for the fear extinction paradigm. Transgenic mice harboring FLEX-L4 conjugating GFP [3] were crossed with homozygous PV specific Cre mice ($PV^{pvr/wt}$) to obtain the mice expressing GFP-L4 specificly in PV interneurons. The room temperature was kept at 23±2°C, and all mice were kept in a room with a 12-hr light/dark cycle (lights on at 6:00 a.m.) with access to food and water *ad libitum*. The female mice used for IntelliCage were group housed (9–12 mice per cage) with limited access to water during the IntelliCage experiments (described below). Adult mice (2 to 5 months old) were used for the behavioral experiments. All experiments were carried out in accordance with the European Communities Council Directive 86/6609/EEC and the guidelines of the Society for Neuroscience and were approved by the County Administrative Board of Southern Finland (License number: ESAVI/38503/2019).

#### Fear extinction paradigm

The fear conditioning paradigm was conducted following a protocol described previously [4]. Briefly, the mice were put into Context A and received an electric foot shock (0.6 mA) after a 30-second sound cue ("beep" sound 80 dB), which was repeated four times with a 30- to 60-second interval (fear conditioning). The mice were equally assigned to two groups: the "control" treated with 0.1% (w/v) saccharine in drinking water and the "experimental group" with 0.1% (w/v) saccharine and 0.080%

(w/v) fluoxetine in drinking water. Two weeks later the mice were put in Context B and received only the sound cue repeatedly for 12 times with different intervals (25-60 seconds) for 2 days (extinction training). One week later, the mice were tested in Context B (spontaneous recovery) followed by exposure to Context A (fear renewal). The duration of the freezing was measured as an index of conditioned fear.

**IntelliCage with chronic fluoxetine treatment**

Intellicage (NewBehavior AG, Zurich, Switzerland) is an automated device that allows housing, performance and measurements of specific tasks in a fully automated manner, removing the need for a human operator and operator-derived bias [5,6]. It consists of a rectangular plastic cage 37.5×55×20.5 cm) with dinking chambers at each of the four corners of the cage. The entrance to each chamber is allowed by a hole with a tubular RFID reader, able to read ID transponders (T-IS 8010 FDX-B, DATAMARS, Switzerland) that are injected subcutaneously in the dorsal-cervical area of each mouse one week before starting the experiment. Each chamber presents two opening doors leading to the nozzles of drinking bottles, and the opening of each door can be controlled electronically through a software. Four red shelters in the middle of the cage provide environmental enrichment and support to reach the food pellet that is provided *ad libitum* on the top of the cage. The set up allows not only the total control of the access to water bottles, but also the recording of the entering in the chamber and nose poking of the door areas leading to the nozzles, made by each mouse, providing an individual behavioural recording for all the animals. Mice were divided into two groups: the “control” treated with 0.1% (w/v) saccharine in the drinking water and the “experimental group” with 0.1% (w/v) saccharin supplemented with 0.08% (w/v) fluoxetine in the drinking water. The mice were gradually introduced to the tasks of the IntelliCage, followed by the leaning tasks. The first two days of housing in the IntelliCage represented the “Free adaptation” phase, during which the animals had full and free access to all corners and water. In the following three days the mice underwent the "Nose poke adaptation" phase, during which they had full access to water only after nose pocking the apertures for the nozzles. In the following four days (Drinking session adaptation, DSA) the mice had to adapt to the specific time window of access to water (21:00-22:00 and 2:00-3:00) used during the Acquisition and Reversal phases. We performed patrolling tasks, where the water bottles were made accessible (doors would open for 4s) only if the mouse nose pocked the "active" door area which, once discovered and used by the mouse, would switch to the one immediately next to it, in a clockwise direction. During the Reversal phase the direction was switched to counter-clockwise. The Acquisition phase of the Patrolling schema would start, lasting for 8 days, followed by 8 days of Reversal.

**Electrophysiology in acute slices**

Mice were deeply anaesthetized with isoflurane and decapitated. The brains were dissected and immersed in ice-cold dissection solution (124 mM NaCl, 3 mM KCl, 1.25 mM $NaH_2PO_4$, 1 mM $MgSO_4$, 26 mM $NaHCO_3$, 15 mM D-glucose, 9 mM $MgSO_4$ and 0.5 mM $CaCl_2$). The cerebellum and anterior part of the brain were removed and coronal 350µm brain slices were cut on a vibratome (Leica Biosystems, Wetzlar, Germany). The slices were incubated for 45 min at 31-32°C in artificial cerebrospinal fluid (ACSF) (124mM NaCl, 3mM KCl, 1.25mM $NaH_2PO_4$, 1mM $MgSO_4$, 26mM $NaHCO_3$, 15mM D-glucose, and 2mM $CaCl_2$) and bubbled with 5% $CO_2$/95% $O_2$. Field excitatory postsynaptic currents (fEPSPs) were recorded in an interface chamber using ACSF-filled electrodes (2-4 MΩ) positioned within the CA1 stratum radiatum. Synaptic responses were evoked every 20 sec. Stimulation intensity was adjusted so that the baseline fEPSP slope was 20-40% of the maximal intensity that resulted in the appearance of a population spike. LTP was induced through tetanic stimulation (200ms pulse interval; 100 pulses; 0.1ms pulse duration) and recorded for 45min.

**Confocal imaging and imaging analysis on PV and PNN intensity**

Immunohistologically stained sections were imaged with a confocal microscope (Zeiss LSM 700). PV/PNN was imaged in different sections containing CA3b region in the dorsal hippocampus (between -1.94 and -2.18 mm in the Anterior-Posterior axis relative to Bregma). Z-Stacks images with 12 -14 layers were obtained with the same acquiring setting to compare the intensity of fluorescence among the samples.

The multilayer confocal images were stacked (z-stack, maximum intensity), and the fluorescence intensity of PV, PNN and TdTomato were analyzed with Fiji software (National Institute of Health, US) (https://fiji.sc/) [7] with the following steps: (1) Each cell was numbered in a stacked image and we measured the size of the area, the mean gray value, the integrated intensity, and the intensity of the surrounding area (mean fluorescence of background readings), (2) the corrected total cell fluorescence (CTCF) was calculated for each cell (*CTCF = Integrated Density - (Area of selected cell x Mean fluorescence of background readings)*. The cells were also classified according to their location in the hippocampal layers (stratum oriens (SO), stratum pyramidale (SP), stratum radiatum (SR)), and cells in only SP were analyzed. We analyzed ten to twenty cells in each image, and three to four images in each group (3-4 animals/group).

**TRAP sample preparation and sequencing**

TRAP-analysis was performed according to a previously published protocol [8]. Briefly, PV-EGFP-L10a mice treated for two weeks with fluoxetine and the non-treated controls (N=3, each group) were euthanized with $CO_2$ directly after the second day of fear extinction. The isolated hippocampi were homogenized and lysed using HEPES –based lysis buffer containing NP-40 (Sigma-Aldrich, Burlington, MA), lipid DHPC (Avanti, Birmingham, UK), and cycloheximide (SigmaAldrich, Espoo, Finland) for stabilizing Ribosome-mRNA complexes, and stored at -80 C until use. Immunoprecipitation of ribosomes binding mRNA was performed with anti-GFP antibody (Memorial Sloan Kettering Cancer Center, New York city, NY) bound to magnetic G-protein coated Dynabeads

(Thermo Fischer, Vantaa, Finland) for 30 min at 4 °C. The beads were precipitated and washed with HEPES –based washing buffer on MagnaRack magnetic stand (Thermo Fischer, Vantaa, Finland). Finally, the mRNA was eluted and purified from the beads using NucleoSpin RNA XS kit (Macherey-Nagel, Germany). Part of the homogenized samples were used as an input for RNA purification to confirm the effect of the immunoprecipitation. The purified RNA was sequenced with HiSeq2500 (Illumina, USA). The sequenced reads were mapped on the mouse genome (GRCm38/mm10), and the number of transcript were counted. Genes with minimal expression were trimmed out and differentially expressed genes were identified using Negative Binomial Distribution –based statistic test with DEseq2 [9]. Enrichment of KEGG-pathways [10] were analyzed for up- and down-regulated genes separately, and significant pathways were detected using Fisher's exact test with DAVID [11]. To narrow down the significant genes, multiple test correction was done using Benjamini-Hochberg method [12].

### Supplemental Figures and tables

Supplemental fig. 1

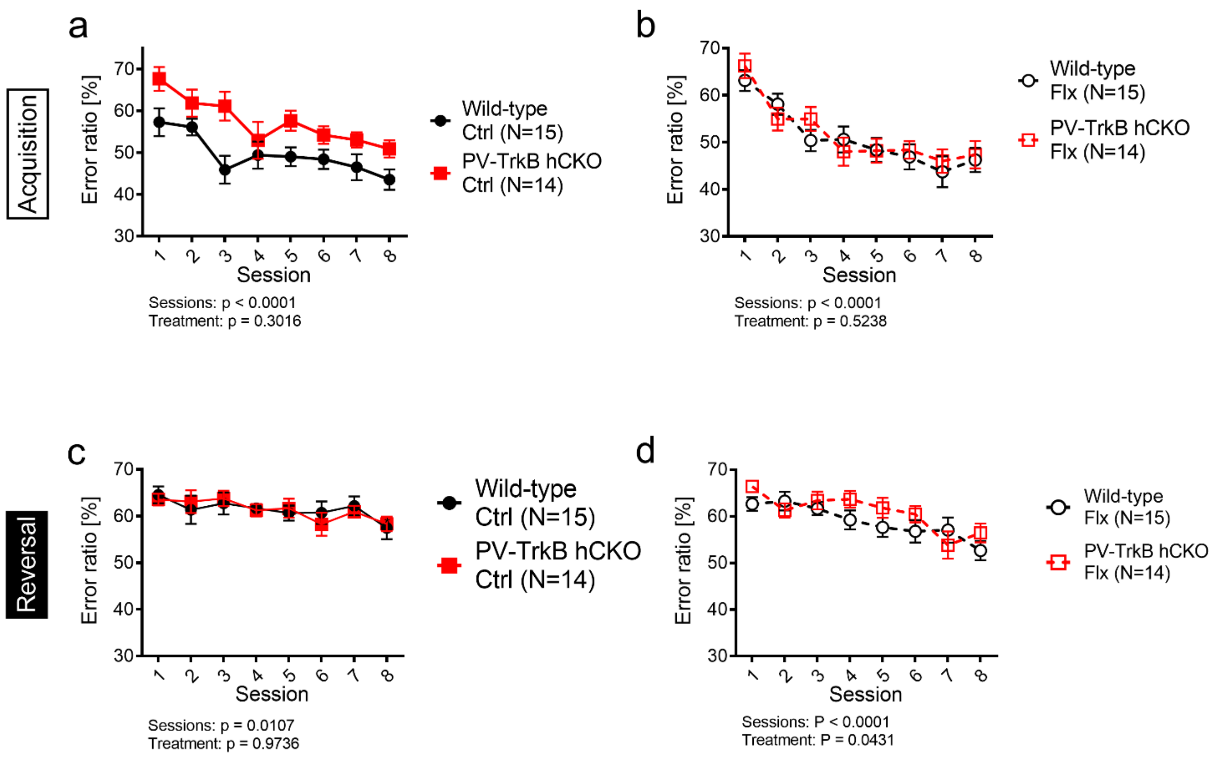


**Supplemental figure 1.**
PV hTrkB cKO mice had a significantly higher error ratio compared to wild-type mice in the control groups($F$ (1,200) = 30.09, $P < 0.0001$) (a), but not in fluoxetine groups ($F$ (1,200) = 0.4079, $P < 0.5238$) (b) in the acquisition phase. There are effects of sessions but no difference in interaction and genotype in both control (c) or fluoxetine treatment (d) in the reversal phase.

**Supplemental table 1. Antibodies used in this study**

| | Host | Company | Product No | Dilution/concentration | Purpose |
|---|---|---|---|---|---|
| anti-GFP antibody | Mouse | MSKCC | HtzGFP_04 (clone19F7) | 178 µg for 1 ml of beads | TRAP |
| anti-GFP antibody | Mouse | MSKCC | HtzGFP_02 (clone 19C8) | 178 µg for 1 ml of beads | TRAP |
| anti-PV antibody | Guinea pig | Synaptic systems | 195004 | 500 | IHC |
| Lectin WFA conjugated with biotin | | Sigma-Aldrich | L1516 | 200 | IHC |
| anti-Guineapig conjugated with Alexa 488 | Donkey | Jackson laboratory | 706-545-148 | 400 | IHC |
| Streptavidin conjugated with Alexa Fluor633 | | ThermoFisher | S21375 | 400 | IHC |

**Supplemental table 2. All detected genes in TRAP analysis (attached excel file)**
Expression of genes in individual samples is represented as ‘counts per million’ (CPM)

**Supplemental table 3. All statistical results (attached excel file)**
